# Transcriptomic impacts and potential routes of detoxification in a lampricide-tolerant teleost exposed to TFM and niclosamide

**DOI:** 10.1101/2023.01.29.526103

**Authors:** M.J. Lawrence, P. Grayson, J.D. Jeffrey, M.F. Docker, C.J. Garroway, J.M. Wilson, R.G. Manzon, M.P. Wilkie, K.M. Jeffries

## Abstract

Sea lamprey (*Petromyzon marinus*) control in the Laurentian Great Lakes of North America often relies on the application of 3-trifluoromethyl-4-nitrophenol (TFM) and niclosamide mixtures to kill larval sea lamprey. Selectivity of TFM against lampreys appears to be due to differential detoxification ability in these jawless fishes compared to bony fishes, particularly teleosts. However, the proximate mechanisms of tolerance to the TFM and niclosamide mixture and the mechanisms of niclosamide toxicity on its own are poorly understood, especially among non-target fishes. Here, we used RNA sequencing to identify specific mRNA transcripts and functional processes that responded to niclosamide or a TFM:niclosamide mixture in bluegill (*Lepomis macrochirus*). Bluegill were exposed to niclosamide or TFM:niclosamide mixture, along with a time-matched control group, and gill and liver tissues were sampled at 6, 12, and 24 h. We summarized the whole-transcriptome patterns through gene ontology (GO) term enrichment and through differential expression of detoxification genes. The niclosamide treatment resulted in an upregulation of several transcripts associated with detoxification (*cyp, ugt, sult, gst*), which may help explain the relatively high detoxification capacity in bluegill. Conversely, the TFM:niclosamide mixture resulted in an enrichment of processes related to arrested cell cycle and growth, and cell death alongside a diverse detoxification gene response. Detoxification of both lampricides likely involves the use of phase I and II biotransformation genes. Our findings strongly suggest that the unusually high tolerance of bluegill to lampricides is due to these animals having an inherently high capacity and flexible detoxification response to such compounds.

## Introduction

The sea lamprey (*Petromyzon marinus*) control program in the Laurentian Great Lakes of North America is one of the most successful invasive species control programs in the world (Siefkes 2017). Sea lamprey control uses lampricide 3-trifluoromethyl-4-nitrophenol (TFM), a highly effective and important control measure (Wilkie et al., 2019). The effects of TFM on non-target fishes are minimised due to the relative inability of lampreys to detoxify TFM (Lech and Statham 1975; Kane et al. 1994; Lawrence et al. 2021). However, TFM sensitivities vary widely amongst non-target fishes found in the Laurentian Great Lakes (Boogaard et al. 2003; Wilkie et al. 2019), with the bluegill (*Lepomis macrochirus*)—a teleost fish belonging to the family Centrarchidae (sunfishes, crappies, and basses), exhibiting one of the highest tolerances to TFM (Boogaard et al., 2003; Lawrence et al., 2021; Wilkie et al., 2019). Indeed, prior work has found that bluegill exhibit a highly diverse set of detoxification genes that likely facilitate high levels of lampricide tolerance (Lawrence and Grayson et al. 2022). Lampricide applications are not restricted to TFM, which is often co-applied with 1-2 % niclosamide (tradename Bayluscide®; previously known as Bayer 73), which increases TFM’s effectiveness without loss of specificity (Barber and Steeves, 2021; Dawson et al., 1982; McDonald and Kolar, 2007). The use of lampricide mixtures is advantageous in control efforts as these two compounds typically act in a synergistic or greater than additive fashion (e.g., Borowiec et al., 2022; Hepditch et al., 2021; Lawrence et al., 2021; Marking and Dawson, 1975), making lampricide applications more effective and economical.

Despite decades of use for sea lamprey control, relatively little is known regarding the physiological impacts of niclosamide on fishes. Historically, niclosamide has been used primarily to treat tapeworm and other parasitic worm infections (Dawson, 2003), by uncoupling oxidative phosphorylation in the target species (Jones, 1979; Pampori et al., 1984; Pearson and Hewlett, 1985; Weinbach and Garbus, 1969). This mode of action has also been shown in vertebrates (Borowiec et al., 2022), which causes increased reliance on anaerobic metabolism to meet basic ATP demands (reviewed in Wilkie et al., 2019; Lawrence et al. 2021; Ionescu et al. 2020a,b). In fishes, this is typified by reductions in tissue ATP, glycogen, glucose, and phosphocreatine levels (PCr; Lawrence et al., 2021; Shoman, 2001; Zhu et al., 2020; Ionescu et al. 2022a,b). Transcriptomic analyses in zebrafish (*Danio rerio*) embryos have shown that niclosamide treatment corresponded with an enrichment of gene ontology (GO) terms associated with cellular growth and metabolism (Vliet et al., 2018; Zhu et al., 2020), phase I and II detoxification (Zhu et al., 2020), as well as physical malformations (Vliet et al., 2018; Zhu et al., 2019). However, our understanding of how non-target species respond to niclosamide exposure and its interaction with TFM remains unclear.

The effects of TFM alone are fairly well characterized as it acts as a mitochondrial uncoupler in fishes (Birceanu et al., 2011; Borowiec et al., 2022; Huerta et al., 2020). Whole transcriptome responses in TFM-exposed bluegill and sea lamprey were characterized by an enrichment of processes related to an arrest of cellular growth and proliferation, increased apoptosis, and an inflammatory response (Lawrence and Grayson et al. 2022). Beyond these works and extensive fundamental toxicological characterizations (Bills and Marking, 1976; Boogaard and Johnson, 2006; Boogaard et al., 2003), the toxicological mode of action and the physiological effects of TFM:niclosamide mixtures in fishes remains underexplored despite the use of TFM:niclosamide mixtures in sea lamprey control in the environment.

In teleost fishes, the primary means of TFM detoxification is thought to involve phase II pathways, in which glucuronidation through UDP-glucuronosyltransferase (UGT) is proposed to be the primary method of TFM conjugation in fishes (reviewed in Wilkie et al., 2019). Indeed, TFM exposure leads to an accumulation of glucuronidated conjugates (Bussy et al., 2018; Lech, 1974; Vue et al., 2002) and upregulation of several *ugt* transcripts (Lawrence and Grayson et al., 2022). Additionally, TFM biotransformation also involves sulfotransferases (phase II) and oxidation through cytochrome P450 (Cyp) proteins (phase I; Bussy et al., 2018; Lawrence and Grayson et al. 2022; reviewed in Wilkie et al., 2019), although less is known regarding the importance of these particular pathways in lampricide detoxification. Due to its phenolic nature, niclosamide detoxification in fishes is predicted to occur through the same general biotransformation pathways as with TFM (Hubert et al., 2005; Statham and Lech, 1975; reviewed in Wilkie et al., 2019). Recent transcriptomic studies on embryonic and larval zebrafish suggest an involvement of both *cyp* and *ugt* genes in teleost (Zhu et al. 2020), as well as mammalian (Lu et al., 2016) niclosamide detoxification.

The present study aimed to identify the transcriptome-wide effects of niclosamide and TFM:niclosamide (1.5%) mixture exposures in bluegill, a tolerant native teleost fish in the Laurentian Great Lakes region. Furthermore, we characterized potential routes of niclosamide biotransformation and elimination. We used a comparative RNA-seq approach to elucidate transcriptome-wide differential expression patterns, a powerful tool for quantifying whole organismal responses (Connon et al., 2018). More specifically, we used gene ontology (GO) term enrichment on differentially expressed genes that responded to the lampricide treatment to summarize the major systems/processes impacted at the cellular level (Gene Ontology Consortium, 2004). Using this approach, we could identify broad scale patterns in cellular function arising from lampricide treatment in bluegill. A GO term enrichment analysis approach has already been used to identify effects on detoxification systems and cellular growth/death processes associated with TFM and niclosamide exposures separately in fishes (Lawrence and Grayson et al., 2022; Zhu et al., 2020). As previous work has identified both TFM and niclosamide as targeting oxidative phosphorylation and affecting cell growth and death processes (Borowiec et al., 2022; Lawrence and Grayson et al., 2022), we hypothesized that both niclosamide and a TFM:niclosamide mixture would disrupt cellular energy metabolism and cell cycle processes in bluegill. We predicted that this effect would be more pronounced in the mixture group. In testing this, bluegill were exposed to niclosamide-alone (0.068 mg L^-1^ nominal), or a TFM:niclosamide (1.5%) mixture (TFM = 4.5 mg L^-1^, niclosamide = 0.068 mg L^-1^ nominal) for up to 24 h. These doses were selected as they represent the concentrations of TFM and niclosamide in the sea lamprey 24 h lethal concentration 99% (LC_99_; concentration needed to kill 99% of the population), which is a close approximation of doses used in field applications of lampricides (Brege et al., 2003). Cellular responses were then inferred using a comparative transcriptomics approach, as such assays are highly useful in characterizing broadscale changes in physiology while also being sensitive enough to determine specific pathways and response patterns (Connon et al., 2018).

## Materials and Methods

### Animal care and collection

The following methodological approach uses the same bluegill exposure series as reported in Lawrence et al. (2021) and Lawrence and Grayson et al. (2022). Herein, we provide a brief overview of our methods, with further details in the aforementioned works. Juvenile bluegill (N = 200; mass = 25.5±0.8 g; total length = 97.3±0.3 mm; Kinmount Fish Farm, Kinmount, ON, Canada) were held at Wilfrid Laurier University in dechlorinated tap water (replacement rate ∼ 0.5 L min^-1^; pH 8.1-8.2, alkalinity ∼255 mg l^-1^ as CaCO_3_, temperature ∼14°C) in 1000 L tanks under a 12 h photoperiod with fish being fed *ad libitum* once daily on commercial feed. All experimental protocols and animal holding were approved by the Wilfrid Laurier Animal Care Committee (AUP # R18001) and followed Canadian Council for Animal Care (CCAC) guidelines.

### Lampricide exposure series

Using field grade TFM (35% active ingredient dissolved in isopropanol; Clariant, Griesheim, Germany) and niclosamide (emulsifiable concentrate containing 16.9% niclosamide ethanolamine salt mixture; Coating Place Inc., Verona, WI, USA), animals were exposed to one of three treatments: a control (i.e., dechlorinated tap water only), a TFM:niclosamide mixture (1.5%), or niclosamide alone, in 14 L glass aquaria (N = 4 bluegill per aquaria) filled with the same aerated water as above (T ∼ 14 °C). The concentration of TFM chosen for the TFM:niclosamide (1.5%) mixture was based on the 24 h LC_99_ of TFM measured in larval sea lamprey in the presence of a 1.5% mixture of TFM:niclosamide (nominal [TFM] = 4.5 mg L^-1^, measured [TFM] = 4.47±0.01 mg L^-1^; nominal [niclosamide] = 0.068 mg L^-1^, measured [niclosamide] = 0.043 ± 0.003 mg L^-1^), which is a close approximation to treatment concentrations used in the field (Brege et al., 2003). The same nominal concentration of niclosamide was used, minus the TFM, for the niclosamide-only exposure (measured = 0.058 ± 0.005 mg L^-1^). Exposures lasted 24 h, with tissue collection at 6, 12, and 24 h. At each sampling time, bluegill were quickly netted, one at time, from the aquaria, and anaesthetized in a bath of buffered tricaine methanesulfonate solution (MS-222; Syndel, Nanaimo, BC, Canada; 1.5 g L^-1^ MS-222 with 3.0 g L^-1^ NaHCO_3_). Tissues (liver, gill) were then collected and immediately transferred to a vial of RNA*later* (Invitrogen, ThermoFisher, Mississauga, ON, Canada) and stored at -80°C for later RNA extraction and transcriptome analyses.

### Extraction of RNA, transcriptome sequencing, and differential expression analyses

Extraction and sequencing protocols are identical to those outlined in Lawrence and Grayson et al. (2022), and they are only briefly outlined here. All extractions were made using a commercial kit (RNeasy Plus Mini Kit; Qiagen, Toronto, ON, Canada) with quality control being determined using a combination of a Nanodrop (260/280 & 260/230 ratios; NanoDrop One Microvolume UV-Vis Spectrophotometer; ThermoFisher, Mississauga, ON), Qubit RNA IQ assay (ThermoFisher, Mississauga, ON, Canada) and a Bioanalyzer (RNA Integrity Number range: 8.5-9.8). Diluted samples (50 ng µL^-1^) were then sent to the Centre d’expertise et de services Génome Québec (Montreal, QC, Canada) for transcriptome sequencing. Here, cDNA libraries were prepared, and then sequenced on a NovaSeq 6000 sequencing system (Illumina, Vancouver, BC, Canada) resulting in 5.13×10^9^ total reads. Raw sequences are available in the National Centre for Biotechnology Information in the Sequence Read Archive (https://www.ncbi.nlm.nih.gov/sra) with the Bioproject accession PRJNA693019.

The resulting libraries were then used to generate a *de novo* bluegill transcriptome. In doing so, we used fish sourced from all three treatment groups and experimental timepoints (6, 12, and 24 h) used here as well as from TFM-exposed bluegill used in complementary studies (see Lawrence and Grayson et al. 2022). The highest read count for a single library from each tissue/timeframe was selected for constructing the assembly (Lawrence and Grayson et al. *2022*) with assemblies performed using Trinity (v2.8.5), and completion then verified using BUSCO (v4.1.4). Following this, the Trinotate (v3.2) annotation protocol (http://trinotate.github.io) was used to annotate resulting Trinity assemblies, which sought to compare open reading frames in the assembly to databases of established proteins, mRNA transcripts, transmembrane helices, signal peptides, and ribosomal RNA sequences, thereby generating likely annotations of our superTranscriptome clusters. Trinity assemblies were then simplified using a Corset-Lace pipeline (https://github.com/Oshlack/Lace/wiki) to hierarchically cluster transcripts creating a final SuperTranscript assembly (Davidson and Oshlack, 2014; Davidson et al., 2017; Langmead and Salzberg, 2012). As a further quality control measure, the resulting superTranscriptome was checked using BUSCO as detailed above. Reads from individual libraries were aligned with the resulting superTranscriptome assemblies using STAR (v2.6.1a) (Dobin et al., 2013), which were used to generate transcript counts via featureCounts (part of subread v2.0.0; Liao et al., 2014).

Resulting count table analyses were conducted in R using R Studio (v1.3.1093; base R v3.5.1; R Core Team (2020)). Here, differential expression of treatment level effects (i.e. lampricide vs control comparison) was made using the package ‘edgeR’ (v3.24.3; McCarthy et al., 2012; Robinson et al., 2010), which employs a quasi-likelihood pipeline. Statistical significance of differentially expressed transcripts was accepted at α = 0.05 with adjustments being made using a Benjamini-Hochberg correction for multiple pairwise comparisons (Kvam et al., 2012). Differentially expressed gene lists (i.e. differences from the controls) were then identified for genes of interest to further our understanding of lampricide toxicity and detoxification. Specifically, we were interested in identifying pathways pertaining to cellular growth and proliferation that have been previously identified as targets of niclosamide including Wnt/β-catenin, Stat-Il6-Jak, Notch, NF-κB, mTor, and Akt signalling (reviewed in Chen et al., 2018). In doing so, we used representative gene targets from within these pathways and, in the case of multiple methods of signalling, elected to use gene targets that are used in the canonical pathways. These search terms included ‘*notch*’, ‘*il6’*, ‘*jak’*, ‘*nfkb’*, ‘*stat’*, ‘*ctnn’*, ‘*axin’*, ‘*wnt’*, ‘*mtor’*, and ‘*akt’* (see Table 1 for gene names). Relevance of the resulting genes from this search were manually curated against UniProt databases (https://www.uniprot.org/). In identifying detoxification genes in the liver and gill of bluegill, we employed the same methodical approach as Lawrence and Grayson et al. (2022) in identifying differentially expressed phase I-III biotransformation genes in the bluegill transcriptome.

**Table 1:** Description of genes associated with growth and proliferation pathways that were searched in the transcriptome of bluegill. Gene identifiers are sourced from the UniProtKB database (UniProt: the universal protein knowledgebase in 2021, 2021).

| Protein Name | Gene ID | Associated Pathway |
| --- | --- | --- |
| Neurogenic locus notch homolog protein | *notch* | Notch Signalling |
| Interleukin-6 | *il6* | Stat-Il6-Jak |
| Tyrosine-protein kinase | *jak* | Stat-Il6-Jak |
| Nuclear factor NF-kappa-B | *nfkb* | NF-κB Signaling |
| Signal transducer and activator of transcription | *stat* | Stat-Il6-Jak |
| Catenin | *ctnn* | Wnt/β-catenin |
| Axin | *axin* | Wnt/β-catenin |
| Protein Wnt | *wnt* | Wnt/β-catenin |
| Serine/threonine-protein kinase mTOR | *mtor* | mTor Signaling |
| RAC-alpha serine/threonine-protein kinase | *akt* | Akt signaling |

This study used enrichment analyses to collate specific pathways and processes, at the transcript level, that were affected by lampricide treatment. Specifically, we characterized GO term enrichment, which identifies important processes associated with genes that were affected by our treatments (Gene Ontology Consortium, 2004). Enrichment analyses were conducted using the R package ‘enrichR’ (v2.1; Jawaid, 2020), allowing us to compare against three databases of interest and derived enriched Gene Ontology (GO) terms in the dataset (https://maayanlab.cloud/Enrichr/; GO_Biological_Process_2018 and GO_Molecular_Function_2018). Here, we focused on two of these databases, including Biological Processes and Molecular Functions, which have an emphasis on higher order collections of molecular systems within overarching biological and molecular level processes, respectively (see Gene Ontology Consortium, 2004 for more in-depth definitions). Resulting GO term lists were then filtered for those that had four or more genes associated with an individual term and were also statistically significant (adjusted p-value < 0.05; Jeffries et al., 2019). The resulting GO term lists were then further summarized using REVIGO (http://revigo.irb.hr/; Supek et al., 2011) at a similarly of 0.5. REVIGO helps to remove redundant GO terms and thus highlight potentially important pathways (Supek et al., 2011). Further details on how these analyses were run can be found in Lawrence and Grayson et al. (2022).

## Results

### Niclosamide transcriptional changes

Niclosamide exposure resulted in differential gene expression in gills, with the peak response occuring at 12 h of niclosamide exposure (47 clusters upregulated, 32 clusters downregulated; Fig. 1A). No genes related to cellular proliferation were differentially expressed (Table 2), but niclosamide treatment resulted in differential expression of several detoxification genes in the gill. Phase I biotransformation genes consisted entirely of *cyp* genes, specifically *cyp1a1* and *cyp1b1*, which were upregulated throughout the niclosamide exposure reaching levels ∼16-fold higher than controls by 24 h (Table 3). Several phase II biotransformation genes were also upregulated and included *ugt3* at all three exposure durations as well as a sulfotransferase (*sult6b1* at 12 h), and three glutathione S-transferases were all upregulated in response to niclosamide treatment (Table 3). Phase III biotransformation genes consisted of a single *abc* that was upregulated (Table 3).

**Table 2:**
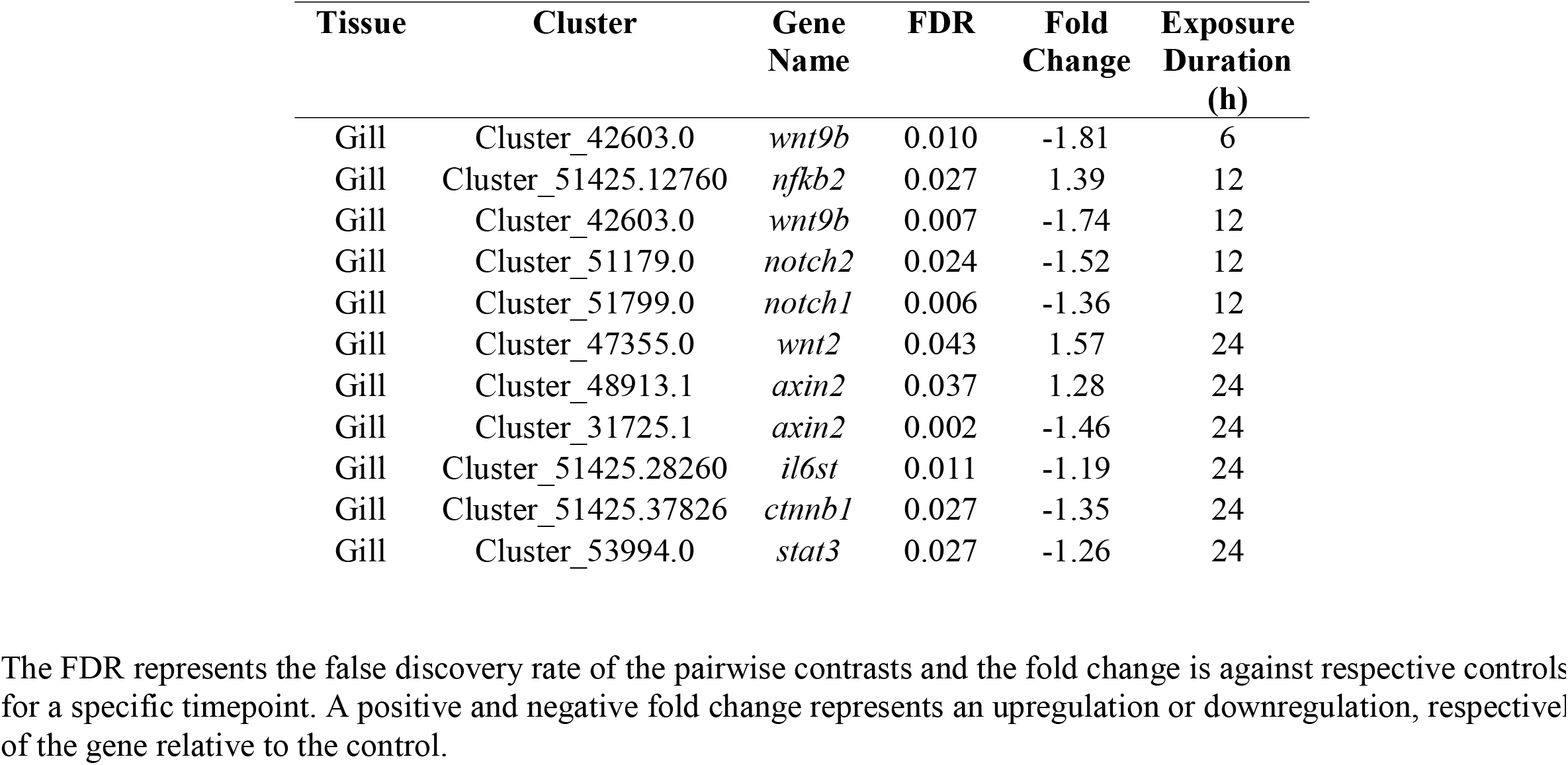
Summary of the differentially expressed gene targets associated with cellular growth/proliferation signalling processes in the gills of bluegill exposed to a TFM:niclosamide (1.5%) mixture (TFM = 4.5 mg L^-1^ nominal, niclosamide = 0.068 mg L^-1^ nominal) for 6, 12, or 24 h. No differential expression of the targeted cellular growth signalling genes was observed in the liver, and niclosamide treatment alone did not result in differential expression of target cellular growth signalling genes in either tissue.

**Table 3:**
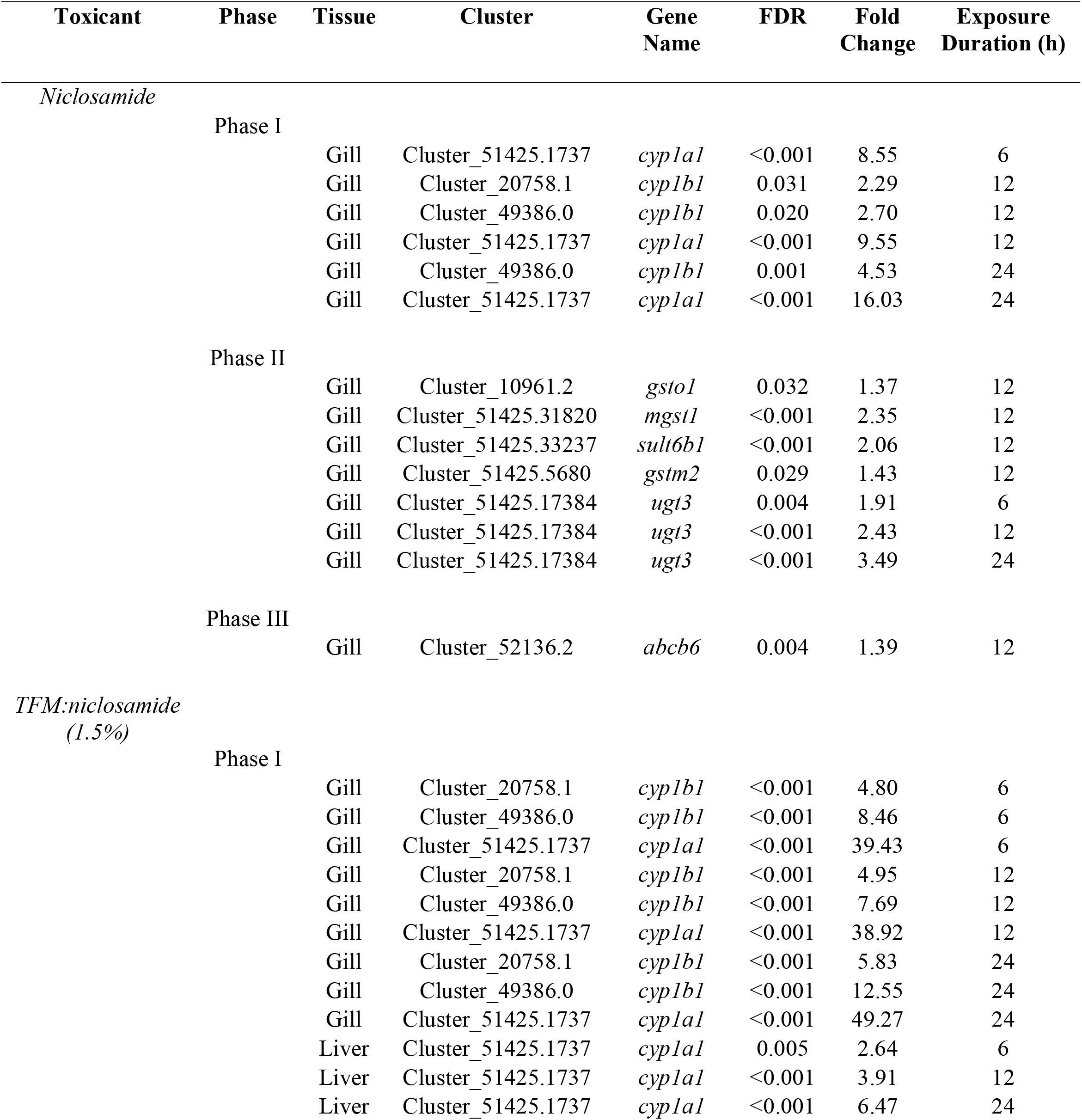
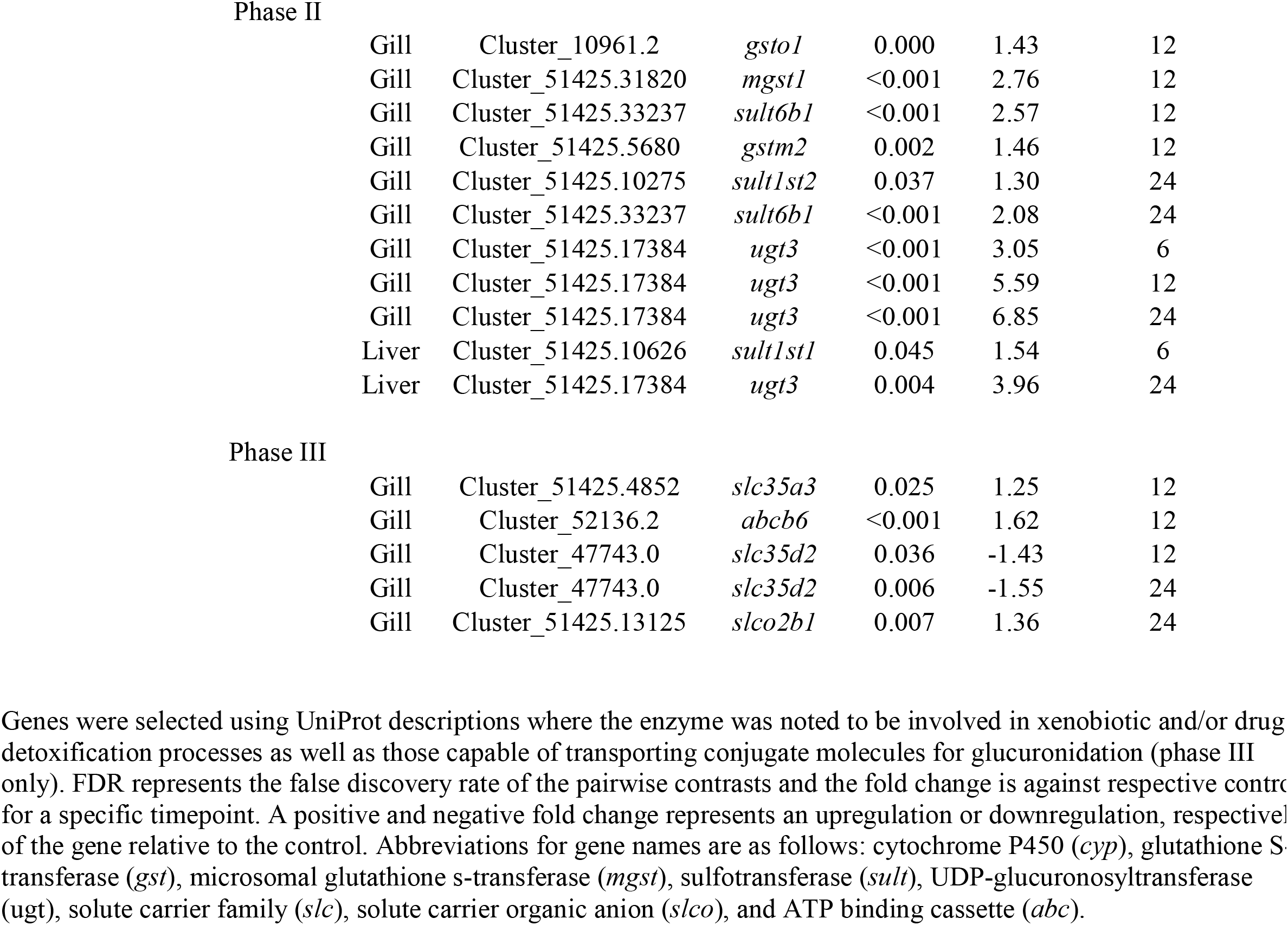
Summary of the differentially expressed genes involved with phase I-III biotransformation processes in the gills and livers of bluegill. Fish were exposed to either niclosamide (0.068 mg L^-1^ nominal) or a TFM:niclosamide (1.5%) mixture (TFM = 4.5 mg L^-1^ nominal, niclosamide = 0.068 mg L^-1^ nominal) for 6, 12, or 24 h.

**Figure 1:**
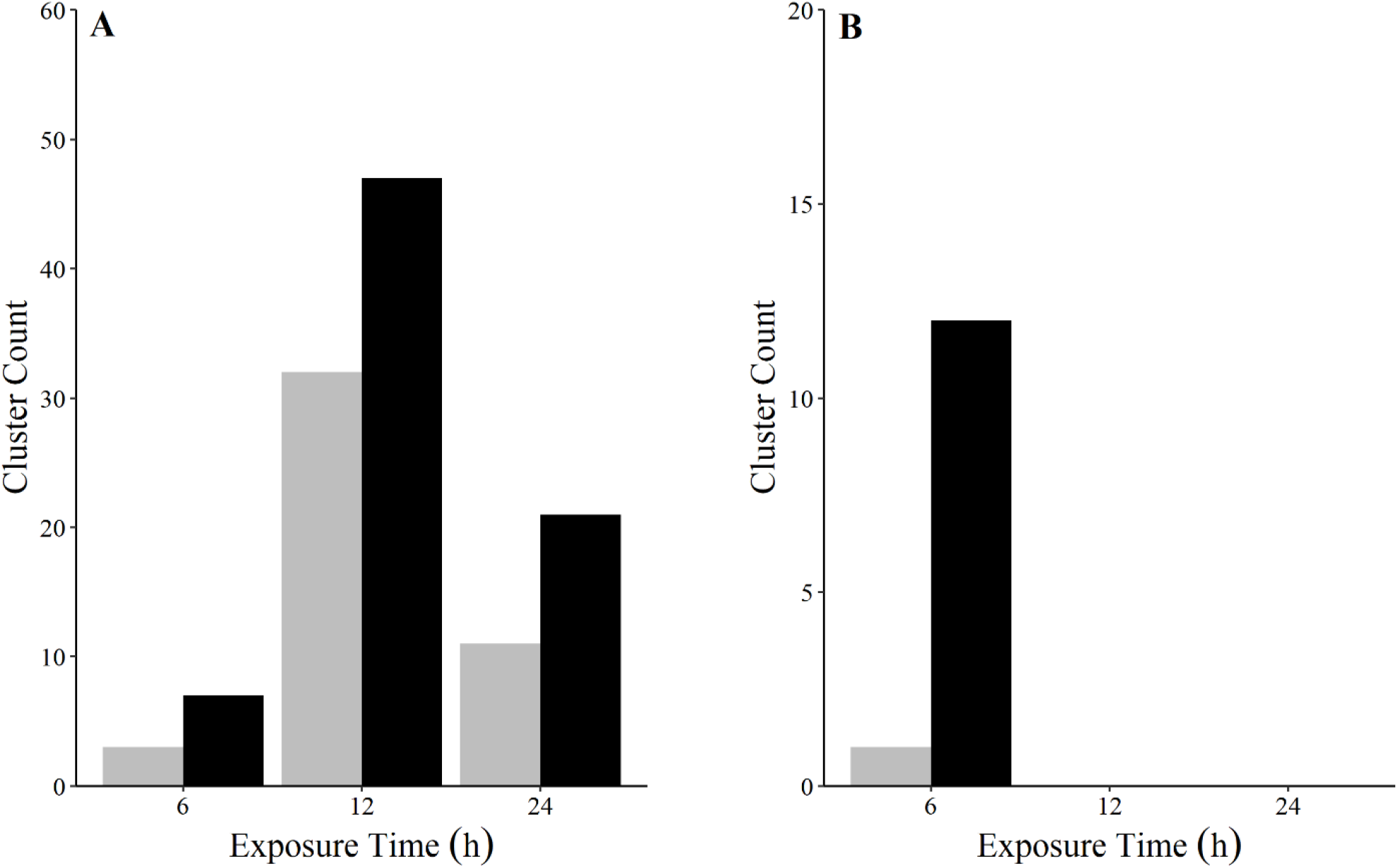
Total counts of differentially expressed superTranscriptome clusters in the gills (A) and liver (B) of bluegill following a 24 h niclosamide exposure (niclosamide = 0.068 mg L^-1^ nominal). Grey bars represent differentially expressed clusters that were downregulated whereas black bars represent upregulated clusters. All clusters were statistically significant at a false discovery rate (FDR) of α = 0.05.

Overall expression patterns in the liver of bluegill were markedly different compared to that of the gill. Differential expression of superTranscriptome clusters was seen only in the liver of bluegill exposed to niclosamide for 6 h, and the overall number of transcripts was lower in liver than in gill (≤ 12 clusters; Fig. 1B). No differential expression of genes associated with either cellular proliferation signalling (Table 2) or biotransformation processes (Table 3) was observed in the liver.

### Mixed lampricide transcriptional changes

Exposure to the TFM:niclosamide (1.5%) mixture resulted in a more pronounced effect on transcription in the gill than niclosamide exposure alone, including increases in the number of downregulated clusters with time, peaking at 490 clusters downregulated by 24 h of exposure (Fig. 2A). Interestingly, clusters associated with upregulated genes reached a maximum of 472 clusters at 12 h of exposure, decreasing thereafter (Fig. 2A). We also identified several gene targets associated with cellular proliferation and growth signalling in the gill, including *wnt9b, notch1*, and *stat3*, wherein most were downregulated in response to the mixture treatment (Table 2). Exposure to the TFM:niclosamide mixture also resulted in a number of biotransformation genes being upregulated (Table 3). For phase I biotransformation, *cyp1a1* and *cyp1b1* were the only genes differentially expressed in the gill, and they were upregulated across all three exposure durations. The *cyp1a1* had the largest change in expression, being ∼39-49-fold greater than controls, compared to *cyp1b1*, which was ∼5-13-fold greater than controls. Differentially expressed phase II biotransformation genes in the gills consisted of an upregulation of *ugt* across all three timepoints (∼3-7-fold higher than controls), as well as higher expression of sulfotransferases and glutathione transferases (Table 3). Unique to the mixture treatment were changes in *slc* expression, with *slc35a3* being upregulated at 12 h of exposure and *slc35d2* expression being downregulated at 12 and 24 h of exposure (Table 3). There was also an upregulation of *abcb6* and *slco2b1* in the gill.

**Figure 2:**
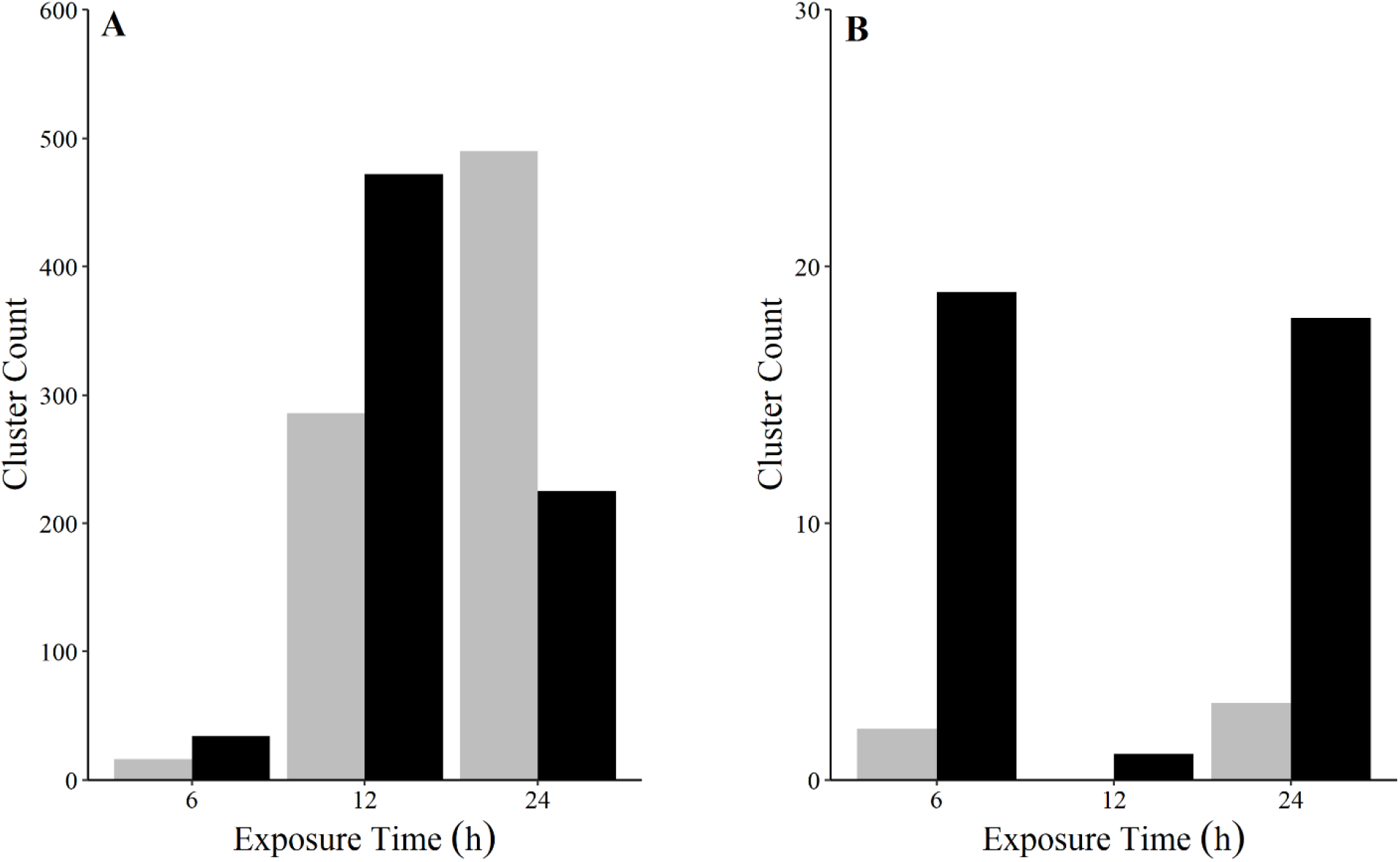
Total counts of differentially expressed superTranscriptome clusters in the gill (A) and liver (B) of bluegill following a 24 h TFM:niclosamide (1.5%) mixture exposure (TFM = 4.5 mg L^-1^, niclosamide = 0.068 mg L^-1^ ; nomin doses). Grey bars represent differentially expressed clusters that were downregulated whereas black bars represent upregulated clusters. All clusters were significant at a false discovery rate (FDR) of α = 0.05.

The liver was less responsive than the gill to the TFM:niclosamide mixture with hepatic superTranscriptome cluster expression showing a more sporadic expression pattern coupled with a lower magnitude of response (Fig. 2B). While downregulated clusters were consistently low throughout the exposure (≤ 3 clusters per timepoint), upregulated clusters were highest at 6 and 24 h with a noticeable drop in transcript abundance at 12 h of exposure (Fig. 2B). No differential expression of cellular growth and proliferation genes were detected in the livers of mixture-exposed bluegill (Table 2). In contrast, biotransformation genes were upregulated in the liver of mixture treated bluegill, which included *cyp1a1*, a sulfotransferase (*sult1st1*), and *ugt3* (Table 3).

### GO term enrichment under lampricide exposure

Most GO term enrichment occurred in the gills of mixture treated fish at 12 and 24 h of exposure. In all, GO term enrichment was dominated by Biological Processes consisting of three main themes in mixture treated fish: influences on cellular growth and proliferation, cell death processes, and immune functions. Unless otherwise noted, results highlight patterns associated with Biological Processes.

Processes associated with cellular growth, proliferation, and survival were the most impacted systems under the TFM/niclosamide mixture treatment. These effects only became apparent by 12 h of the mixture treatment and continued through to the 24 h exposure duration. For upregulated genes at 12 h of exposure, ‘DNA replication’ constituted the largest enrichment component (Fig. 3). Within this parent term, we observed several daughter terms that suggest the cell cycle processes were impacted by the mixture treatment as ‘DNA replication’, ‘DNA metabolic process’, ‘negative regulation of transcription, DNA templated’, and ‘anaphase-promoting complex-dependent catabolic process’ were all enriched (Fig. 3). With respect to Molecular Functions, upregulated genes at 12 h also corresponded with an enrichment of cyclin protein kinase activities, further suggesting that the cell cycle was affected by the TFM:niclosamide (1.5%) mixture (Fig. 4). Larger impacts on cell growth and proliferation were observed in 12 h downregulated genes. Here, ‘regulation of MAPK cascade’ corresponded with a downregulation in several processes associated with cellular proliferation/growth (Fig. 5). Stem cell differentiation was also downregulated here further suggesting that the mixture treatment is repressing cell growth while promoting apoptosis. With respect to Molecular Functions, there was an enrichment of the growth and proliferation activities associated with phosphatidylinositol 3-kinase activity (Fig. 6).

**Figure 3:**
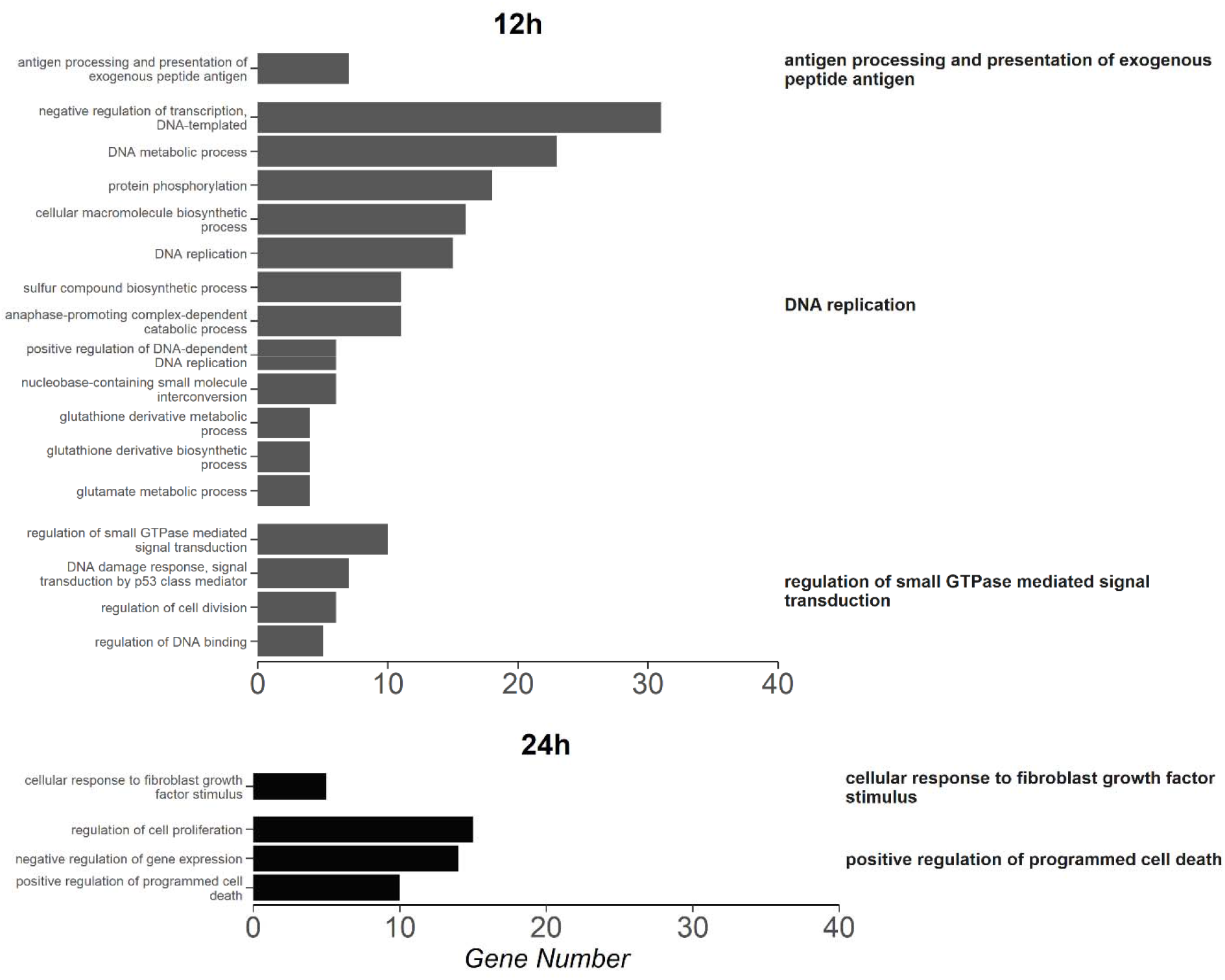
Summary of enriched gene ontology (GO) terms, corresponding with Biological Processes, in genes that we upregulated following 12 h (dark grey) and 24 h (black) of TFM:niclosamide (1.5%) mixture (3-trifluoromethyl-4’-nitrophenol; TFM = 4.5 mg L^-1^ & niclosamide = 0.068 mg L^-1^; nominal doses) exposure in the gills of bluegill (*Lepom macrochirus*). Genes were considered differentially regulated at a false discovery rate < 0.05. Only GO terms from the functional analysis with an adjusted p < 0.05 and at least four transcripts were considered significantly enriched. REVIGO was used to summarize GO terms to reduce redundancy and group according to similarity (right labels). Par terms are presented on the right-hand side of the figure while child terms associated with each of these are to their immediate left.

**Figure 4:**
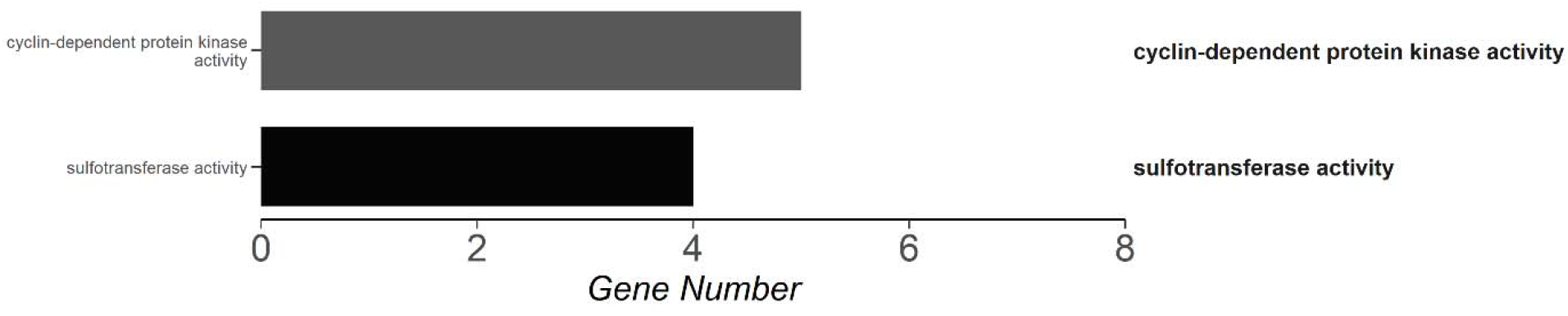
Summary of enriched gene ontology (GO) terms corresponding with Molecular Functions in genes that wer upregulated following 12 h (dark grey) and 24 h (black) of TFM:niclosamide (1.5%) mixture (3-trifluoromethyl-4’-nitrophenol; TFM = 4.5 mg L^-1^ & niclosamide = 0.068 mg L^-1^; nominal doses) exposure in the gills of bluegill (*Lepom macrochirus*). Genes were considered differentially regulated at a false discovery rate < 0.05. Only GO terms from the functional analysis with an adjusted p < 0.05 and at least four transcripts were considered significantly enriched. REVIGO was used to summarize GO terms to reduce redundancy and group according to similarity (right labels). Par terms are presented on the right-hand side of the figure while child terms associated with each of these are to their immediate left.

**Figure 5:**
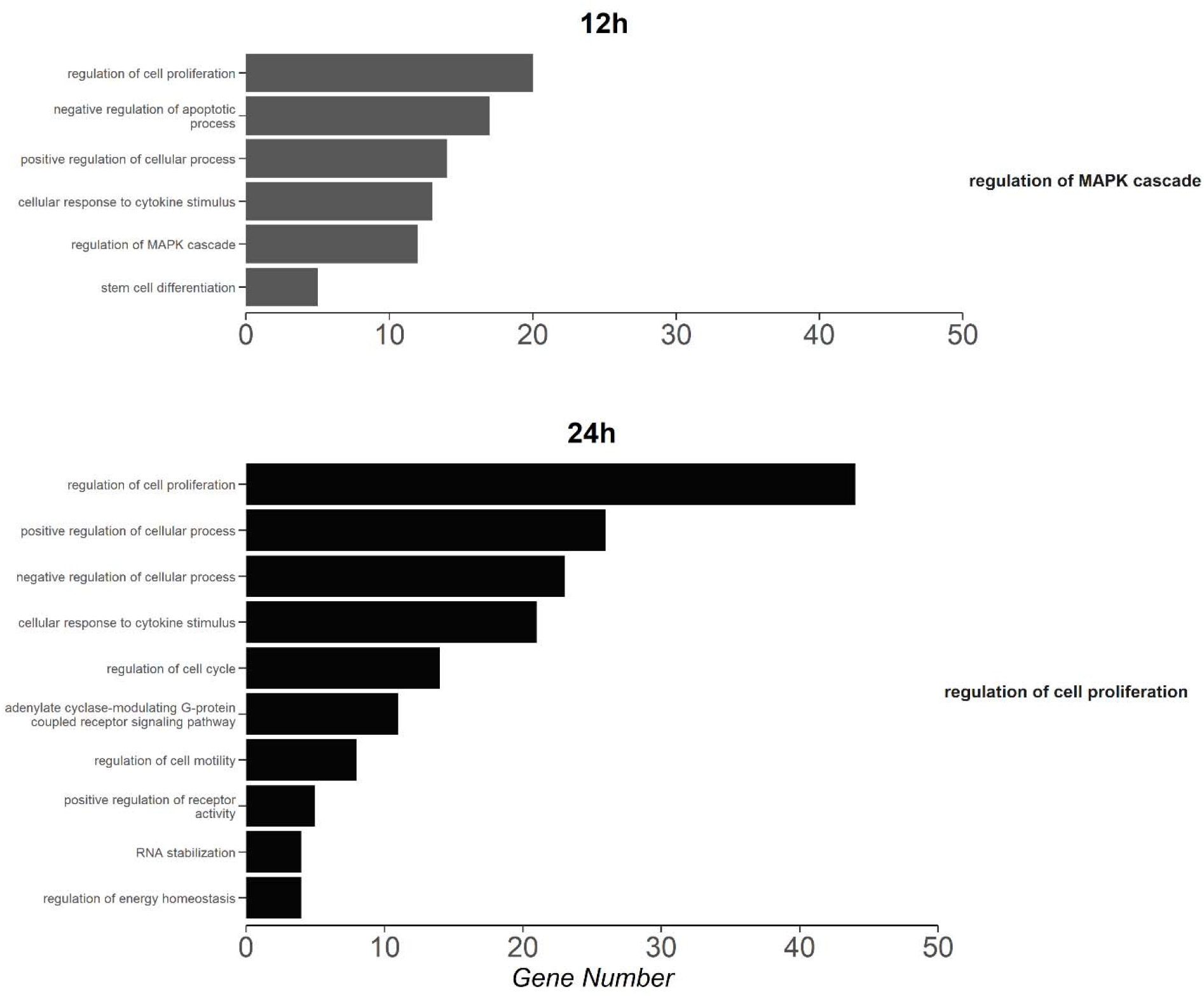
Summary of enriched gene ontology (GO) terms, corresponding with Biological Processes, in genes that we downregulated following 12 h (dark grey) and 24 h (black) of TFM:niclosamide (1.5%) mixture (3-trifluoromethyl-4’-nitrophenol; TFM = 4.5 mg L^-1^ & niclosamide = 0.068 mg L^-1^; nominal doses) exposure in the gills of bluegill (*Lepom macrochirus*). Genes were considered differentially regulated at a false discovery rate < 0.05. Only GO terms from the functional analysis with an adjusted p < 0.05 and at least four transcripts were considered significantly enriched. REVIGO was used to summarize GO terms to reduce redundancy and group according to similarity (right labels). Par terms are presented on the right-hand side of the figure while child terms associated with each of these are to their immediate left.

**Figure 6:**
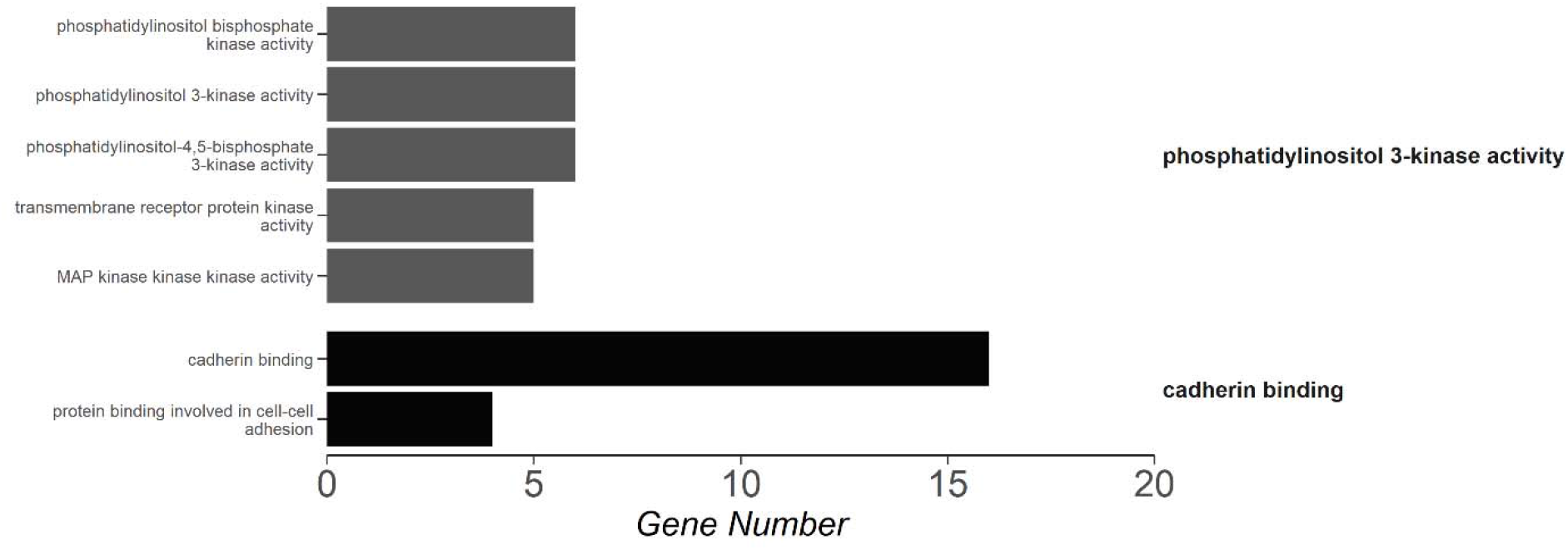
Summary of enriched gene ontology (GO) terms corresponding with Molecular Functions in genes that wer downregulated following 12 h (dark grey) and 24 h (black) of TFM:niclosamide (1.5%) mixture (3-trifluoromethyl-4’-nitrophenol; TFM = 4.5 mg L^-1^ & niclosamide = 0.068 mg L^-1^; nominal doses) exposure in the gills of bluegill (*Lepom macrochirus*). Genes were considered differentially regulated at a false discovery rate < 0.05. Only GO terms from the functional analysis with an adjusted p < 0.05 and at least four transcripts were considered significantly enriched. REVIGO was used to summarize GO terms to reduce redundancy and group according to similarity (right labels). Par terms are presented on the right-hand side of the figure while child terms associated with each of these are to their immediate left.

By 24 h of exposure, the gill was still exhibiting enrichment of GO terms associated with cell growth, the cell cycle, and apoptosis. For genes that were upregulated, there was a large contribution of GO terms that were associated with apoptosis (parent term: ‘positive regulation of programmed cell death’; Fig. 3). Concurrently, there was also an enrichment of GO terms associated with ‘regulation of cell proliferation’ in upregulated genes.

Downregulated genes at 24 h of exposure were enriched for genes encompassed by the parent GO term ‘regulation of cell proliferation’ (Fig. 5). Within this parent term, the ‘regulation of cell proliferation’ was the largest and was accompanied by enrichment of ‘regulation of the cell cycle’. Interestingly, there was both positive and negative regulation of cellular process observed here (Fig. 5). Tangential to some of the growth processes outlined here, exposure to the TFM:niclosamide mixture appeared to have an impact on cellular structural elements, as ‘cadherin binding’ was found to be enriched in downregulated genes at 24 h of exposure for Molecular Functions (Fig. 6).

Exposure of bluegill to the TFM:niclosamide mixture also had a significant influence on expression of genes associated with immune response. In upregulated genes at 12 h of exposure, mixture treated fish had an enrichment of GO terms associated with antigen processing (Fig. 3). For downregulated genes at 12 h of exposure, there did appear to be some impacts of immune function-related genes as demonstrated by the enrichment of the ‘cellular response to a cytokine stimulus’ (Fig. 5). By 24 h of exposure, immune responses were more evident. Indeed, mixture-exposed gills had an enrichment of GO terms associated with ‘cellular response to fibroblast growth factor stimulus’ in upregulated genes by 24 h (Fig. 3). In downregulated genes, cytokine responses were also observed at 24 h (Fig. 5).

## Discussion

We used a transcriptomics approach to characterize the broader modes of toxicity of niclosamide and a TFM:niclosamide mixture, while also elucidating potential lampricide detoxification pathways in a non-target teleost fish. By evaluating the suites of genes that were up- and downregulated in the gills and liver of bluegill within 6–24 hours of exposure, we were able to provide insights into the molecular and biological processes involved in the response of this relatively lampricide-tolerant fish to these chemicals. In line with our predictions, exposure to the TFM:niclosamide mixture had a greater impact on the transcriptome of bluegill than niclosamide alone. While niclosamide had almost no effect, the lampricide mixture was associated with an enrichment of GO terms related to cell cycle arrest and growth. As evident in the transcriptomic profiles, mixed lampricide detoxification likely involves a diversity of phase I and phase II biotransformation and phase III detoxification, which is thought to lead to greater tolerance in bluegill (Lawrence and Grayson et al. 2022). Overall, we provide novel characterizations of the toxicological responses to niclosamide and a TFM:niclosamide mixture in a non-target teleost fish while also inferring potential routes of lampricide detoxification. As sea lamprey control efforts are increasingly aiming to minimize impacts on native, non-target species (Siefkes et al., 2013), integrating a more comprehensive understanding of the potential adverse effects and routes of elimination of lampricides is required in developing more informed and targeted approaches to sea lamprey control.

### The associated toxicological impacts of a TFM:niclosamide mixture

Exposure to a TFM:niclosamide mixture involved an enrichment of processes related to an arrest of the cell cycle, and growth and cellular death. While these are the first transcriptomic data examining niclosamide-TFM interactions in a teleost, our results align with the effects of these toxicants on their own. For example, both niclosamide (Jin et al., 2010; Liu et al., 2015; Wieland et al., 2013) and other phenolic compounds similar to TFM, notably 2,4-dinitrophenol (Han et al., 2008a; Han et al., 2008b; Lam et al., 2013; Miyoshi et al., 2006), are capable of arresting the cell cycle, limiting cellular proliferation, and enhancing apoptosis in a clinical setting. In fishes, TFM itself has been shown to enrich GO terms related to cell death and arrested growth in both bluegill and sea lamprey (Lawrence and Grayson et al. 2022). Together these observations suggest a common mechanism of toxicity for TFM and niclosamide (Wilkie et al. 2019). Interestingly, transcripts associated with cell growth and death were localized to the gills, which may be explained by exposure to high environmental concentrations of the lampricides compared to the liver exposure levels (see Lawrence et al., 2021) as well as potentially having lower constitutive expression of detoxification enzymes, relative to the liver (Christen and Fent, 2014; Husoy et al., 1994; Santovito et al., 2012).

### Potential routes of TFM:niclosamide detoxification

Mixture-exposed bluegill had a greater diversity and magnitude of effect (i.e., higher fold changes) for detoxification genes relative to niclosamide exposure alone. In fishes, TFM detoxification is dominated by glucuronidation and, to a lesser extent, sulfation (Bussy et al., 2018; Hunn and Allen, 1975; Lech, 1974; Lech and Costrini, 1972; Vue et al., 2002). Similar patterns have also been observed in invertebrates (Kawatski and Bittner, 1975) and mammals (Lech, 1971). In line with this, our results appear to demonstrate Ugt’s as the main biotransformation enzymes involved in a transcriptomic response to the mixture as it features in both niclosamide (Zhu et al., 2020) and TFM (Bussy et al., 2018; Lech, 1974) detoxification. Furthermore, *ugt3* was the most responsive of the phase II biotransformation genes (fold changes ≤ 6.85x); an effect observed previously with TFM exposures in bluegill (Lawrence and Grayson et al., 2022). The Cyp’s role in biotransformation is limited given that TFM’s structure already facilitates phase II conjugation (Wilkie et al. 2019), although prior work identified possible Cyp-mediated TFM metabolite formation in both teleosts and sea lamprey (Bussy et al. 2018a,b). Further kinetic studies and metabolite characterizations would be required to confirm the involvement of Cyp’s in TFM biotransformation and the specific processes by which *ugt3* functions in lampricide detoxification.

In contrast to niclosamide exposures, we also observed differential expression of *slc* and *abc* genes following mixture treatment. As these proteins generally function in the transport of toxicants, conjugates, or detoxification substrates, they may function in either an excretory role or in limiting environmental uptake of lampricides in the gill (Stott et al., 2015). The large amount of branchial gene expression is likely the product of this tissue being directly exposed to high concentrations of lampricide (see Lawrence et al., 2021). As suggested by Lawrence and Grayson et al. (2022), the gill may function to limit the uptake of lampricides from the environment (i.e., increased efflux and decreased influx) while also functioning in phase I and II biotransformation thereby limiting the parent form of the lampricide in circulation. While our data does not explore either of these possibilities, flux studies could be useful in determining the fate and movement of lampricides across the branchial epithelium and could help in addressing some of these possibilities.

### Niclosamide toxicity and potential detoxification pathways

Bluegill have been previously reported to show little physiological disturbance under niclosamide treatments (i.e., 24 h at 0.05 mg L^-1^; Lawrence et al., 2021), and we also identified little differential gene expression under niclosamide-only exposures. This null effect may result from the low exposure dose as the 24 h LC_50_ for niclosamide in centrarchids is more than 0.09 mg L^-1^ (Marking and Hogan 1967). Likely, the high detoxification capacity (see below) coupled with the low exposure dosage posed little challenge to bluegill resulting in a minimal transcriptomic signature of niclosamide toxicity.

Early work with mammals and invertebrates identified reduction, hydrolysis, and glucuronidation as being important in niclosamide metabolism (Douch and Gahagan, 1977; Griffiths and Facchini, 1979; Kawatski and Zittel, 1977). In fishes, glucuronidated and sulphated niclosamide metabolites are present in the bile and muscle tissues (Dawson et al., 1999; Hubert et al., 2005; Statham and Lech, 1975). These observations, coupled with the accessible hydroxyl group on niclosamide (reviewed in Smart and Hodgson, 2018), have implicated Ugt and Sult proteins in niclosamide biotransformation (Wilkie et al., 2019). Indeed, Ugt-mediated niclosamide glucuronidation has been directly observed in isolated rat liver and intestinal microsomes incubated with UDP-glucuronic acid (Fan et al., 2019). Additionally, five species of phenolic compounds, which may be considered analogous to niclosamide, were almost entirely metabolized through glucuronidation and sulfation in zebrafish (Kasokat et al., 1987). Accordingly, we observed upregulation of both *sult6b1* and *ugt3* in the gill of bluegill. At the transcriptome level, this has also been observed previously as Zhu et al. (2020) found that chronic niclosamide exposure (120 h at 0.04 mg L^-1^) produced an upregulation of *ugt* genes in zebrafish. However, our results provide support for phase II biotransformation in niclosamide detoxification, that not only involves Ugt1 and Ugt2 enzymes, but also Ugt3.

Higher *cyp* expression levels under niclosamide treatment are not likely important in niclosamide detoxification here. Functionally, Cyp proteins increase water solubility via hydroxylation and oxidation reactions (reviewed in Smart and Hodgson, 2018). Given that niclosamide already has an accessible hydroxyl group, it is unlikely that Cyp proteins have a strong involvement in niclosamide detoxification and that simply employing phase II metabolism is sufficient for eliminating the toxicant. However, Bussy et al. (2018a) demonstrated that TFM generated a reactive amino metabolite, likely via Cyp450 mediated oxidation. While *cyp* genes are upregulated in niclosamide invertebrates (Zhang et al. 2015; Buddenborg et al., 2019; Fan et al. 2019) and fishes (Zhu et al. 2020) previously, functional characterizations of niclosamide metabolites have failed to detect Cyp-mediated niclosamide catalysis in fishes (Stathem and Lech 1975; Allen et al. 1979; Dawson et al. 1996, 1999). Interestingly, while Cyp-mediated hydroxylation of niclosamide has been observed in isolated mammalian liver microsomes (Lu et al. 2016; Fan et al. 2019), its *in vivo* contribution remains minimal, with Ugt-mediated metabolism being the dominant detoxification pathway (Fan et al. 2019). Furthermore, Cyp1a2, which was not found differentially expressed in bluegill here, had the highest contribution to niclosamide hydroxylation with Cyp1a1 and Cyp1b1 having near zero contributions (Lu et al. 2016). Likely, differential expression of *cyp* genes here contribute to another niclosamide-responsive physiological process, perhaps being involved in energy metabolism or a stress response element. It is also possible that there was sufficient Cyp protein in the gill (i.e., high constitutive expression) such that differential expression changes were not needed. This would be especially true if more preferable detoxification systems such as Ugt or Sult have sufficient capacity to detoxify lampricides. We may expect to see changes in *cyp* expression if these systems become saturated (i.e., operating at V_max)_ and alternative detoxification systems become necessary.

## Acknowledgements

The authors would like to thank Oana Birceanu for logistical assistance, Fisheries and Oceans Canada for providing the TFM used in this study, Darren Foubister and Josh Sutherby for conducting the exposures, Kimberly Ta for mRNA extractions, Sara Good and Matthew Doering for their helpful discussion and comments related to the de novo assembly and the gene annotations, and the crew at the U.S. Geological Survey’s Hammond Bay Biological Station for collecting the larval lamprey used in this study.

## Author Information

KMJ, MPW, RGM, JMW, CJG, and MFD were responsible for the experimental design. The labs of KMJ and MPW carried out the exposures and sample collection. Data and sample analyses were conducted by MJL, PG, and JDJ. The first draft of the manuscript was written by MJL and PG, with all authors contributing to the refinement and production of the final manuscript.

